# Predicting tumour mutational burden from histopathological images using multiscale deep learning

**DOI:** 10.1101/2020.06.15.153379

**Authors:** Mika S Jain, Tarik F Massoud

## Abstract

Tumour mutational burden (TMB) is an important biomarker for predicting response to immunotherapy in cancer patients. Gold-standard measurement of TMB is performed using whole exome sequencing (WES), which is not available at most hospitals owing to its high cost, operational complexity, and long turnover times. We developed a machine learning algorithm, Image2TMB, which can predict TMB from readily available lung adenocarcinoma histopathological images. Image2TMB integrates the predictions of three deep learning models that operate at different resolution scales (5X, 10X, and 20X magnification) to determine if the TMB of a cancer is high or low. On a held-out set of patients, Image2TMB achieves an area under the precision recall curve of 0.92, an average precision of 0.89, and has the predictive power of a targeted sequencing panel of approximately 100 genes. This study demonstrates that it is possible to infer genomic features from histopathology images, and potentially opens avenues for exploring genotype-phenotype relationships.

## INTRODUCTION

Immune checkpoint inhibitors have induced significant and long-lasting responses in patients with non-small cell lung cancer (NSCLC)^1–3^. Pembrolizumab is FDA approved for first-line treatment of metastatic NSCLC in combination with chemotherapy, and as monotherapy in patients with high *PD-L1* (programmed death-ligand 1) expression. Nivolumab is FDA approved for second-line treatment of metastatic NSCLC. Despite remarkable responses in many patients, early investigations have revealed that not all patients benefit from immunotherapies. While *PD-L1* expression is a biomarker of response to immunotherapy, a significant number of patients with low *PD-L1* expression and only half of patients with high *PD-L1* expression benefit from pembrolizumab^1^. The search for novel predictors of response to immunotherapy is therefore essential.

Tumour mutational burden (TMB) is emerging as an important biomarker for predicting response to immunotherapy^4–6^. TMB is a measure of the total number of nonsynonymous somatic mutations per megabase of the tumour genome coding area^3^. Tumours with high TMB are thought to express a greater diversity of neoantigens, resulting in increased immune recognition when immune checkpoint inhibitors release natural brakes on the immune system^3^. Several studies have shown that response to immunotherapy is associated with high TMB in patients with advanced solid tumours^3,6,7^. In addition, using high TMB as an indication for combination nivolumab/ipilimumab therapy has shown promise in early clinical trials^8^. Oncologists are therefore increasingly considering TMB in their decision to prescribe immunotherapy. Low cost, reliable, and fast assays for TMB are critically needed.

Whole exome sequencing (WES) is the gold-standard for measurement of TMB. While feasible in research settings in which the utility of TMB has been determined, WES is not currently used in the clinical oncology arena owing to prohibitively high costs and logistical constraints, including lengthy turnaround times^9^. However, clinicians now routinely use lower-cost next generation sequencing of targeted panels of genes. These targeted panels guide use of gene-directed therapies and entry into clinical trials, with a typical turnaround time of two to four weeks. Efforts are underway in clinical laboratories to repurpose these targeted sequencing assays to predict TMB that is determined when using WES, by using the default prediction technique of normalizing the number of mutations found by the covered sequencing area^10^. However, major hurdles for use of panel sequencing for TMB prediction include inadequate sampling owing to the small fraction of the exome sequenced, and the targeting of genes that are recurrently mutated in cancer, which introduces bias into the normalized TMB estimate^10^. For these reasons, larger panel sizes, at minimum 1-3 megabases, and a paired-normal sample are needed for robust normalized TMB estimates^11^. This requires adding sequencing space for many genes, including non-recurrently mutated genes for which clinical utility other than TMB may not be established. Larger panel sizes also increase cost and force important trade-offs between sequencing depth and number of patients per sequencing run. In most clinical settings, these logistical constraints potentially increase turnaround times beyond the usual two to four weeks for targeted panel sequencing. Development of an alternative simpler method to assess TMB would therefore be enormously beneficial.

While histopathologists have long recognized the association of certain cancer morphological phenotypes with mutations in individual genes, TMB has not been considered in that regard to date. We hypothesized that aggregated mutations across individual tumour cells result in global morphological changes that are detectable in routine histopathological images when using machine learning techniques. Machine learning employing handcrafted features on histopathological images^12,13^, has previously been used to differentiate subtypes^14^, and predict survival outcomes^15^. Recently, deep learning has been applied to biomedical images and promises more accurate and robust predictions^16^. Deep convolutional networks have been applied to histopathological images to detect tumours^17,26^, differentiate subtypes of NSCLC, and predict certain gene mutations in NSCLC^18,19^.

Herein, we develop a deep learning method, named Image2TMB, for predicting TMB in lung adenocarcinoma (LUAD) by using digitized images of frozen hematoxylin and eosin (H&E)-stained histopathological slides from 499 patients in The Cancer Genome Atlas (TGCA) (https://www.cancer.gov/tcga). We focus on developing an approach for LUAD owing to the availability of both image and sequence data for a large number of patients with this malignancy as compared to other cancer types in the TGCA, and owing to the particular success of TMB as a biomarker in the treatment of lung cancer^5,8^. Image2TMB uses Inception v3^20^, a convolutional neural network (CNN) that has achieved state-of-the-art performance on the ImageNet Large Scale Visual Recognition Challenge^21^. Image2TMB classifies patients as high TMB with good performance, including within important subgroups of patients stratified by smoking status and whether they contain pathogenic variants for *TP53* (tumour protein p53), *EGFR* (epidermal growth factor) and *KRAS* (Ki-ras2 Kirsten rat sarcoma viral oncogene homolog). We believe that with further development and clinical validation, our method offers a potential alternative assay to determine TMB with at-diagnosis turnaround times and very low cost. In addition, Image2TMB demonstrates that biological images contain features that represent nucleotide changes in the whole genome, and not just in a few genes. Our results indicate that histopathological images contain previously unexplored yet clinically useful features that are detectable using deep learning techniques.

## RESULTS

An overview of Image2TMB and the dataset we used is outlined in Fig. 1, and the performance of Image2TMB is shown in Fig. 2. Image2TMB achieved good performance on an independent test set of patients, with an AUC of 0.92 and the average precision (AP) of 0.89 (Fig. 2a). Interestingly, the multi-scale model aggregating across the three magnifications performed substantially better than any of the scales on their own. The best single-scale performance was achieved by the 20X magnification, with an AUC of 0.81 and AP of 0.74; higher magnification was generally better for prediction than lower magnification. The improvements from multi-scale aggregation suggested that there was complementary information that the deep learning model can capture from the different levels of granularity in the images. As an additional validation, we applied Image2TMB to slides of non-cancerous lung tissue biopsied from the test patients that were not used during the training of our algorithm (Fig. 2b). For patients with a TMB above 206, probability predictions were significantly lower for these slides as compared to the cancer slides. Probability predictions of non-cancerous slides did not significantly differ between the patients above and below a TMB of 206, unlike predictions of the cancer slides. This provides evidence that Image2TMB was utilizing features unique to cancerous tissue to make predictions, and correctly predicted non-cancerous tissue having a low TMB even when the patient had a cancer with a high TMB.

**Figure 1.**
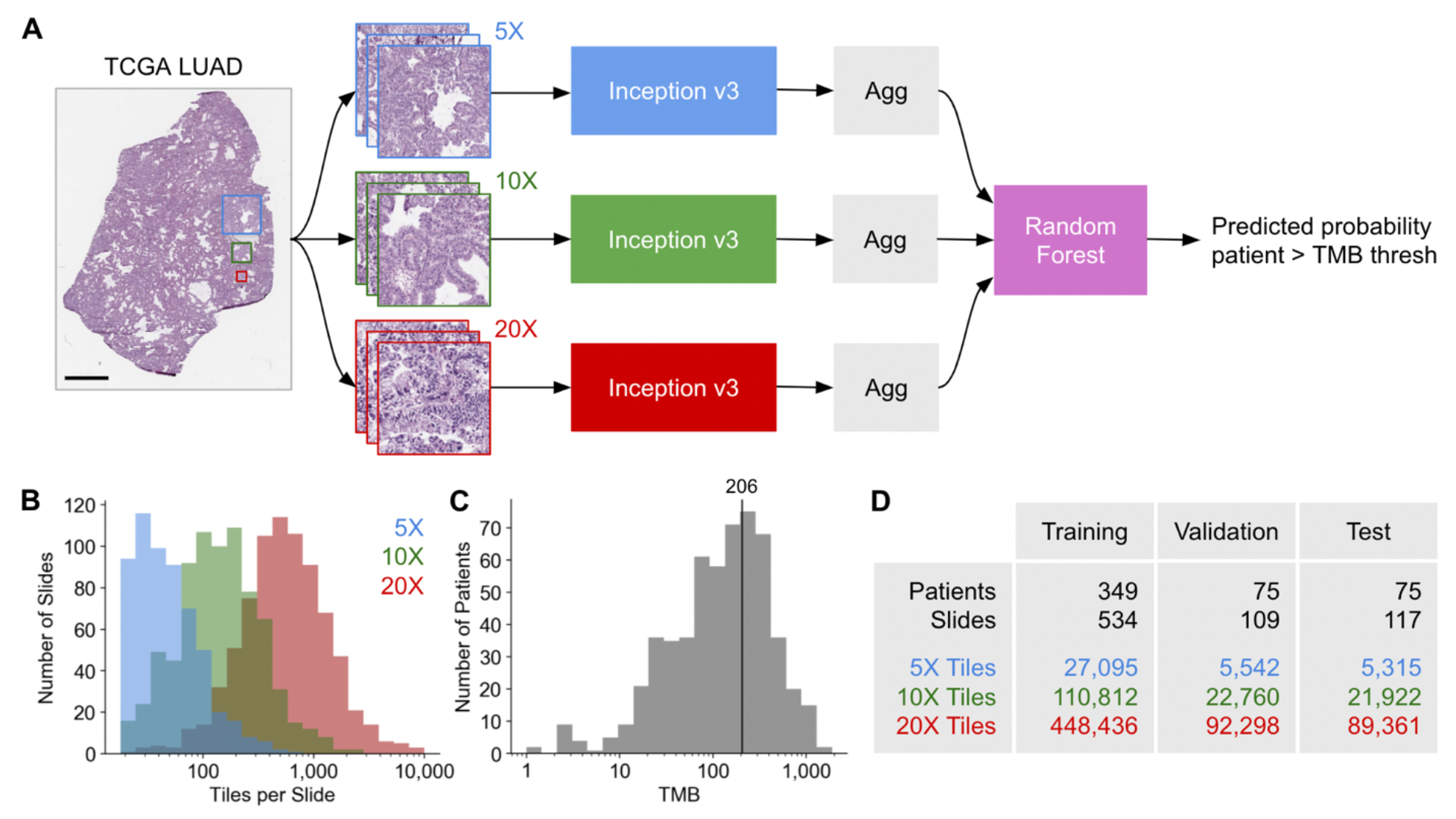
Overview of Image2TMB and dataset. **a**, Computational pipeline of Image2TMB. A given whole-slide image is divided into tiles with 5X, 10X, and 20X magnifications. Convolutional neural networks based on the Inception v3 architecture predict the probability of high TMB for each tile. The tile-level predictions are aggregated (Agg) into a single prediction at each magnification by removing predictions between 0.3 and 0.7 and then taking the median. A random forest uses the aggregated results from the three magnifications to predict whether the TMB of the patient is above or below a given threshold. Scale bar indicates 1 mm. **b**, Histogram of the number of tiles per slide. **c**, Histogram of TMB across all available cases in the TCGA LUAD dataset. Line indicates the threshold used to define high and low TMB. **d**, Number of patients, slides, and tiles in the training, validation, and test datasets.

**Figure 2.**
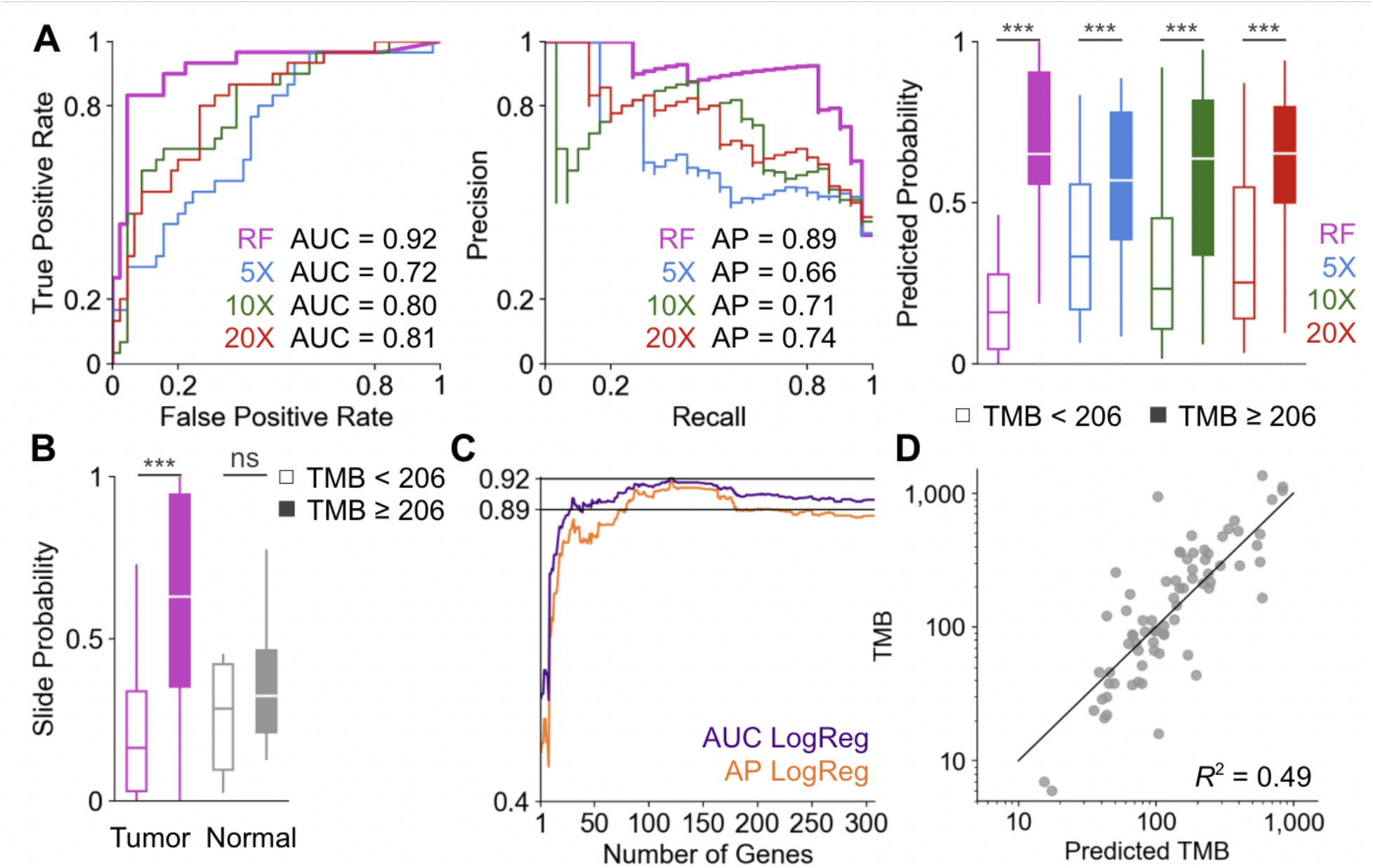
Performance of Image2TMB. **a**, Area under the receiver operator curves (AUC, left), area under the precision recall curves (middle), and box plots of the patient probability predictions of the Image2TMB and aggregated predictions at each magnification (right). **b**, Slide-level probability predictions of slides with tumour tissue versus slides with tissue uninvolved by tumour. **c**, The area under the ROC curve (AUC) and the average precision (AP) of the random forest is compared against a baseline model consisting of a logistic regressor (LogReg) over a panel of selected genes. **d**, Scatter plot of TMB regression. Confidence intervals are in Supplementary Table 1. All results are reported only for patients in the test dataset.

We chose the binary classification task of predicting TMB above and below a given threshold because TMB was being tested as an indication for immunotherapy. The same Image2TMB model could also be adapted to predict the TMB number itself as a regression task, and it achieved an R^2^ of 0.49 on the test patients (Fig. 2d).

To place Image2TMB in context, we compared its performance with that of predicting TMB from targeted sequencing of gene panels, which is common in clinical practice^9^. We selected the gene panel with a greedy procedure: we started with the single gene whose gene-level TMB was the most highly correlated with the exome TMB, and then iteratively added genes that maximally improved the performance of the panel-based predictor. The panel predictor was a logistic regression that predicted the exome TMB from the TMB of the individual genes in the panel. In the clinical setting, targeted sequencing panel genes are not selected based on their TMB predictive values and these clinical gene panels would likely perform worse than the panel results we report here for a similar number of genes. Fig. 2c shows the AUC and AP of the gene panel as its size increases. The performance of Image2TMB was comparable to panels of 100 genes. The advantage of Image2TMB is that it can be immediately obtained from the readily available histopathological images, while targeted sequencing of 100 genes typically requires two to four weeks and costs several hundreds of dollars to achieve.

Next, we examined the performance of Image2TMB on different subgroups of patients. We stratified the patients by their smoking status and whether they contained pathogenic variants for *TP53, EGFR* and *KRAS,* which are frequent in lung cancer. *TP53* and *KRAS* mutations have been associated with high TMB, while *EGFR* mutations have been associated with low TMB^22–24^. Within every subgroup of patients, smokers and non-smokers, and wild type and carrier for variants in these three genes, Image2TMB consistently achieved high performance (Fig. 3). This further validated the robustness of the method when applied to different scenarios.

**Figure 3.**
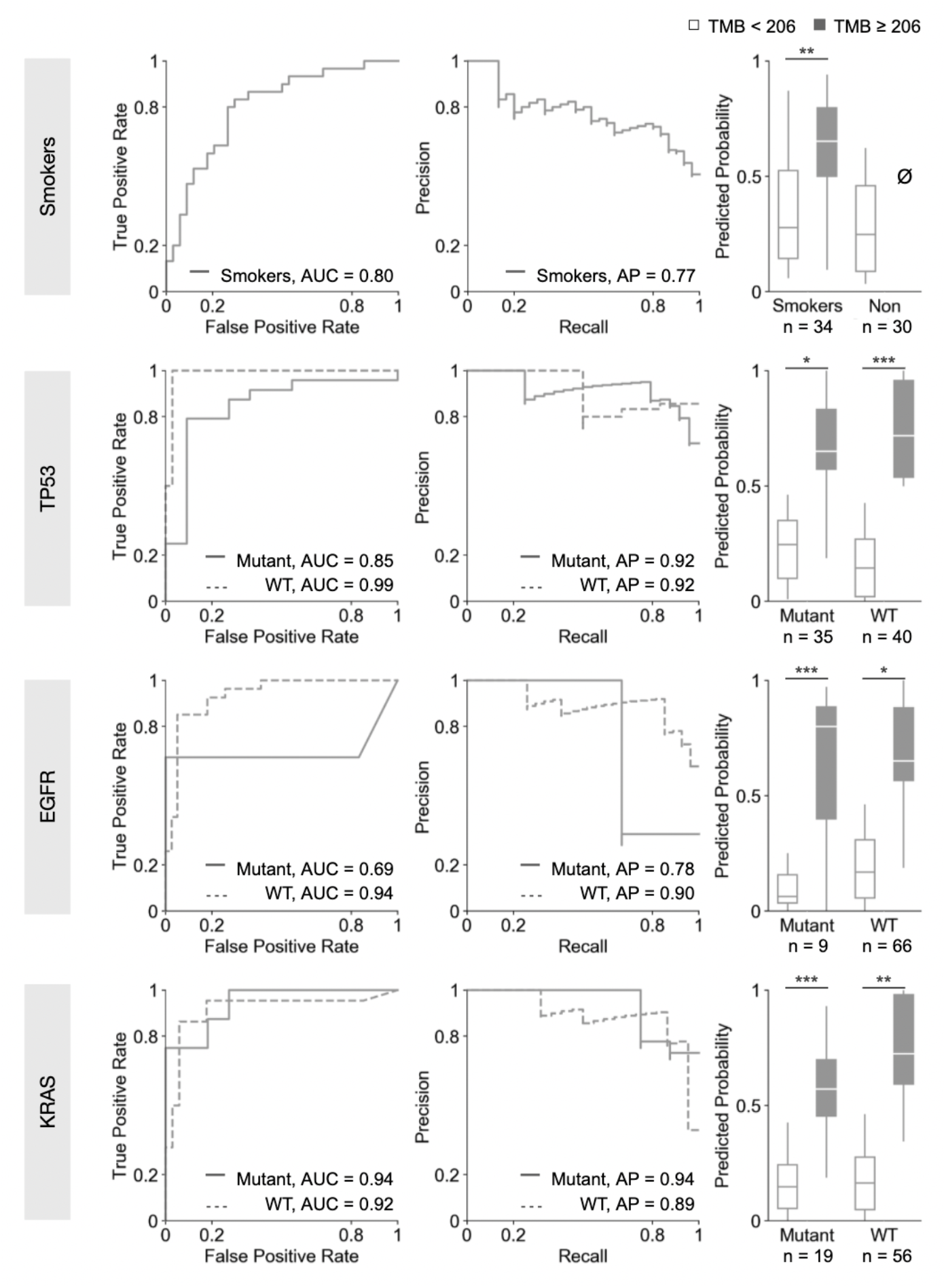
Stratification study of TMB prediction. Receiver operator curves, precision recall curves, and box plots of probability predictions in patient populations stratified by smoking history (smoker or nonsmoker) and whether a pathogenic mutation was detected in *TP53, KRAS,* and *EGFR.* WT is wild type. All non-smokers have TMBs below 206. All results are reported only for patients in the test dataset.

To validate the robustness of Image2TMB, we trained it to predict other TMB thresholds of 135.5, 223, and 293.5, which correspond to the median, upper tertile, and upper quartile of TMBs of patients available in the TCGA LUAD dataset. We found that the AUC, AP, comparison to selected panels of genes, calibration, regression, performance on different patient subgroups, and Matthew’s Correlation Coefficient (MCC) were all comparable across all thresholds (Supplementary Fig. 1-3, Supplementary Table 2).

Image2TMB also provides a visualisation of the spatial distribution of mutational burden predictions. The tile-level prediction of Image2TMB can be used to generate heatmaps of the spatial TMB predictions at three different resolutions for each histopathological image (Fig. 4). Red areas of the heatmap indicated regions of the tissue predicted by the algorithm to have high TMB. Predicted regions of high TMB were consistent across the different magnifications, even though the models at different magnifications were trained independently. Image2TMB predicted regions of both high and low TMB within the bounds of tumours, suggesting the model may be detecting morphological correlates of TMB that are heterogeneous across the cancer.

**Figure 4.**
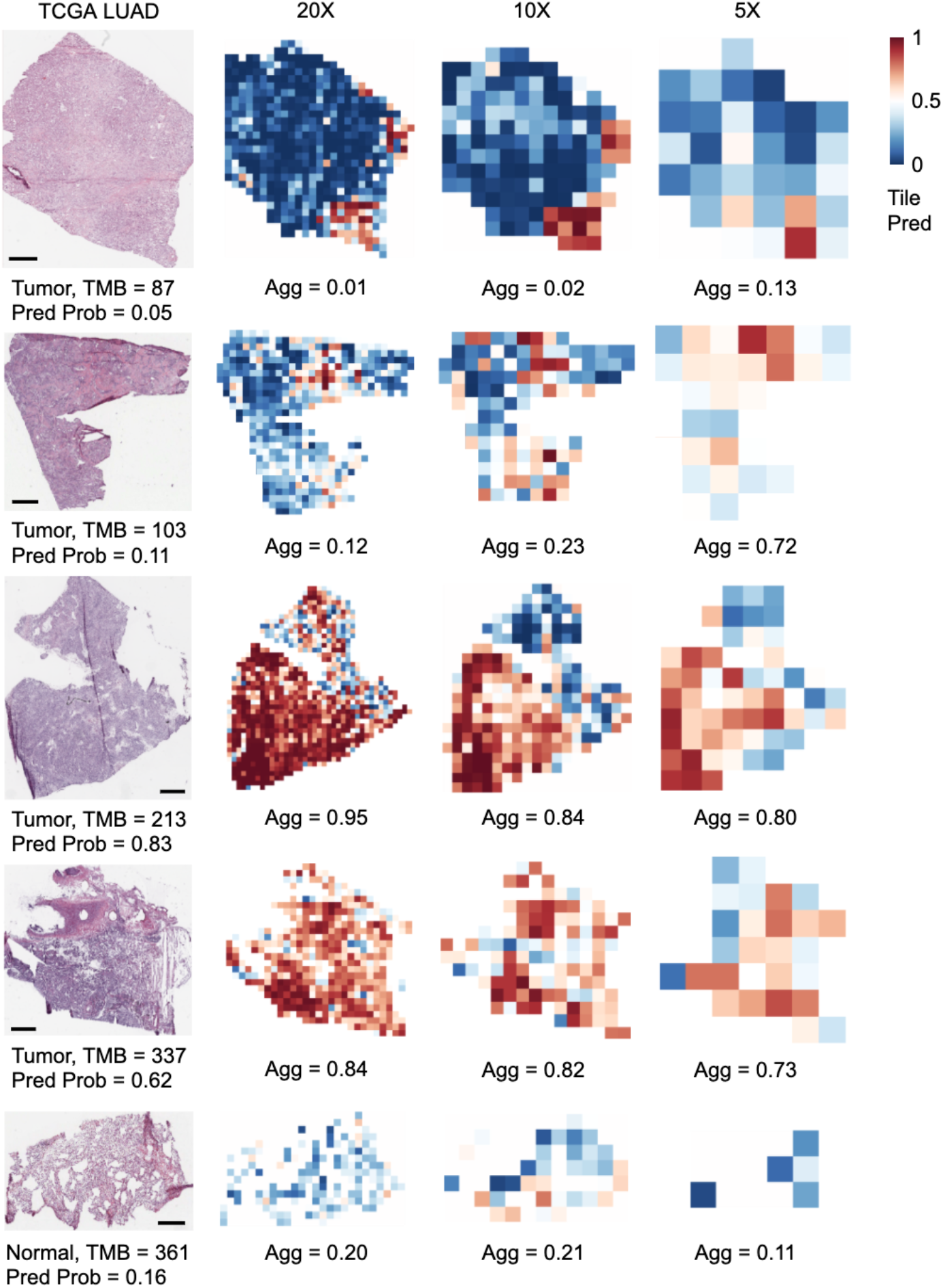
Spatial heterogeneity of predictions. For five random patients in the test set, we show a raw histopathological slide and heatmaps of the tile predictions at each magnification. The bottom slide is lung tissue that is uninvolved by tumour (from a patient with a tumour with TMB = 361), while the other slides show tissues involved by tumour.

## DISCUSSION

TMB is emerging as an important correlate of response to immunotherapy and a possible indication for immunotherapy in NSCLC patients. Here, we demonstrate that TMB can be assessed in LUAD using digitized frozen-section histopathological images with similar classification ability to targeted sequencing of 100 genes covering approximately 300 kilobases. While thresholds for immunotherapy treatment indications have not been fully determined, our technique demonstrates good performance across a wide range of thresholds, showing robustness of Image2TMB to threshold choice.

Targeted sequencing for prediction of TMB is currently constrained by high cost and lengthy turnaround times. In addition, the specific genes included in panels vary between institutions, resulting in differing biases in estimation of TMB when using targeted next generation sequencing. In contrast, the method we present here is limited only by computational and slide digitization logistical and cost considerations. Compared to targeted sequencing that can take two to four weeks to accomplish, it takes approximately 2–2.5 min to scan histopathological slides in the same manner as was used for TCGA, and less than 1 min to run our entire pipeline on the histopathological slides of a single patient. Once scanned, the same histopathological slides used by pathologists for diagnosis potentially could be used to predict TMB.

Currently, the generalisability of our method is limited by the training set. In clinical practice, patients with advanced stage malignancy are likely to have relatively small diagnostic biopsy specimens (rather than surgical resections), which may be from one of many different potential sites of involvement including lymph nodes, pleural fluid, liver and other locations. To enable practical applications of our approach, training and testing in these different scenarios is necessary. In addition, the TCGA frozen-section images used to train the convolutional neural network are highly enriched for cancer cells, which does not reflect real-world tissue samples acquired at the time of biopsy or surgery. Histopathologists, however, could manually define cancerous regions within slide images, and apply Image2TMB on only those regions of interest. Frozen-section slides can often be of poor quality, with tissue fold artefacts, varying staining, and differing section thickness. Despite this, our method shows good performance, suggesting that this method is robust to variance in specimen processing. Given these promising results using frozen-section slides, future studies using formalin-fixed paraffin-embedded tissue slides, which are of superior quality, may be successful. As the same stains and similar staining techniques are used across institutions, we believe that with further development and a larger training set, the method presented here could be generalizable and standardised.

Further steps are needed to validate Image2TMB in a clinical setting. Further clinical validation would include comprehensive comparison to alternatives with different model architectures and hyperparameters, evaluation of the response of Image2TMB to heterogeneity in LUAD images and genomes, cohort studies to best choose the TMB threshold, and ultimately, clinical trials to study immunotherapy outcomes alongside such an approach.

Our method shows performance in classification similar to that of large targeted sequencing panels. However, targeted sequencing panels have superior performance in quantitative prediction, with panels covering greater than one megabase demonstrating an *R^2^* greater than 95%^10^. Reasons for this large discrepancy in classification and regression performance can likely be attributed to the fact that our deep learning models were trained only to determine if TMB was above or below a threshold, not to predict the exact TMB value. Targeted sequencing panels display poor quantitative predictive performance and increased variance in specimens with low TMB, as gene panels only sample up to a hundred out of 19,000 to 20,000 genes in the human genome^10^. Because our method does not rely on gene sampling, a notable advantage of our method is that prediction error is likely to be unaffected by low TMB levels. This is consistent with our results, which show low variance in prediction at low TMB levels compared to targeted sequencing panels (Fig 2d).

Our work finds that TMB, a clinically relevant measurement that conventionally requires additional and laborious testing, can be predicted from the H&E stained histopathological images in the TCGA LUAD dataset. Additionally, in contrast to other diagnostic tools, such as exome sequencing and panels, Image2TMB predicts the TMB for each region of the image. This may represent heterogeneity of histologic features associated with high TMB, or alternatively could represent heterogeneity in TMB itself. In contrast to current approaches for bulk measurement of TMB, Image2TMB shows that deep learning could assist in the study of the spatial heterogeneity of tumours. Image2TMB is also an initial step in applying deep learning to connect histopathological images with a whole-genome-level features, which would aid in the study of relationships between cancer genotypes and phenotypes. Image2TMB is therefore a step towards enhancing the usefulness of the vast quantities of readily available histopathological images, to help screen and prioritise patient samples and subsequent treatments.

## Acknowledgements

The results presented are based upon data generated by the TCGA Research Network: https://www.cancer.gov/tcga. We would like to thank R. Joshi, N. Neishaboori, and E. Nohr for feedback and comments on the paper.

## Author contributions

M.J.S. conceived and performed the study and experiments. T.F.M. supervised the experiments. M.J.S. and T.F.M. wrote the manuscript. Both authors discussed the results and commented on the manuscript.

## Competing interests

The authors declare no competing interests.

## Additional information

Supplementary information is available for this paper at …

Reprints and permissions information is available at www.nature.com/reprints.

Correspondence and requests for materials should be addressed to M.J.S. or T.F.M.

## Data availability

All data used in this study are publicly available via the TCGA Research Network: https://www.cancer.gov/tcga.

## Code availability

Our full approach, including data download, preprocessing, and Image2TMB, is publicly available from Github: https://github.com/msj3/Image2TMB.

## METHODS

### Data processing

We used whole-slide histopathological images and whole exome sequences from The Cancer Genome Atlas (TCGA), which is a collaboration between the National Cancer Institute (NCI) and the National Human Genome Research Institute (NHGRI). We used all available LUAD cases for this study, consisting of 499 patients associated with 760 H&E stained whole-slide images of frozen tissue sections. Our computational approach, Image2TMB, and the TCGA dataset are summarized in Fig. 1. Because the whole-slide images were many thousands of pixels in each dimension, they were too large to be used as direct input to a neural network. We instead divided the whole-slide images into 5X, 10X, and 20X magnification tiles that were all 512 × 512 pixels. This resulted in tens to thousands of tiles per slide (Fig. 1c). The whole-slide images were randomly split into three sets: training, validation and testing datasets, with 349, 75, and 75 patients, respectively (Fig. 1b). Each patient and their associated slides were assigned to only one of these train, validation, and test datasets, ensuring that there was no overlap between the datasets. We used the training and validation datasets for all of the development and training of our computational models, while the test dataset was used only to report final results. Crucially, this guarantees that our model was never trained and evaluated on the same tiles, slides, or patients. We took the TMB of each patient to be the total number of nonsynonymous mutations in the whole exome sequence of the patient’s cancer. Mutations were identified with the VarScan2 variant caller^25^.

### Deep learning on histopathological images

Image2TMB first made TMB predictions separately for every tile across the three magnifications. We used deep neural networks based on the Inception v3 architecture to predict the probability of high TMB for each tile. A separate Inception v3 model was trained for each magnification level. Inception v3 is a CNN that uses inception modules consisting of several convolution operations performed in parallel with different kernel sizes and one max pooling layer. We combined five convolution nodes with two max pooling operations and followed this by eleven stacks of inception modules. The last component of the architecture was a fully connected layer with a softmax operation. This produced two normalized probability predictions corresponding to the probability the tile was from a patient who has a TMB above or below a particular TMB threshold. We focused on a TMB of 206 as the target threshold because this threshold approximates 10 mutations per megabase when using genes found in the FoundationOne CDX panel, and has been used in clinical trials previously^8^. In our dataset, 38% of patients had a TMB above 206, making this a relatively balanced prediction task. To validate the robustness of our model, we also trained the same architecture to predict other TMB thresholds of 135.5, 223, and 293.5, which correspond to the median, upper tertile, and upper quartile of TMBs of patients available in the TCGA LUAD dataset (Supplementary Fig. 1-3). We initialized our network parameters with pre-trained weights from the ImageNet competition. We then trained all model parameters using backpropagation on the tiles in the training dataset. The loss function was defined as the cross entropy between predicted probability and the true class labels.

### Patient-level prediction

Image2TMB then aggregated the tile probabilities to produce a single probability per patient per magnification. This was accomplished by removing low confidence tile predictions (probabilities between 0.3 and 0.7) and then taking the median of all predictions for a given patient and magnification. This produced three probabilities per patient each corresponding to a different magnification. A random forest (RF) classifier was then trained on the validation dataset to predict from these three probabilities if a patient’s TMB was above the threshold. This prediction by the RF was the final, patient-level prediction of our approach. The probability that a patient had high TMB (i.e. above the threshold) was taken to be the proportion of votes of the trees in the ensemble. RFs were separately trained for each threshold using only the validation dataset. It is this integration across magnifications to produce a single prediction that makes our approach multiscale.

### Performance evaluation

After training, the performance was evaluated and reported using the test dataset. We derived receiver operator and precision recall statistics at several TMB thresholds of the patient-level prediction and the predictions of all Inception v3 models. We estimated confidence intervals by 10,000 iterations of the bootstrap method. Patient-level accuracy was compared against a baseline model consisting of logistic regression over a limited panel of genes that predicts if a patient has a TMB over a particular threshold. The logistic regressor was trained on patient vector with each element corresponding to a gene, and with a value of one if the gene has a nonsynonymous mutation, and zero otherwise. The logistic regressor was fit to the training dataset and we examined its area under the curve and average precision on the test dataset. We increased the number of genes in the panel by adding the gene that most increased classification accuracy on the validation dataset. For patient subset analysis, smoking status was determined using information provided by TCGA. Pathogenicity of mutations in *TP53, KRAS,* and *EGFR* was determined by review of current literature and annotation databases, including OncoKB and My Cancer Genome.

### Hyperparameter and model selection

All hyperparameter and model selection was conducted based on performance on the validation dataset. This included the selection of the multiscale approach, use of Inception v3 model, use of the RF, optimization parameters, and the 0.3 and 0.7 thresholds for filtering the tile prediction. The test dataset was used only after our approach was fully developed, and to report all results presented in this research. No modifications were made to our approach after evaluation with the test dataset. Crucially, this ensured we did not violate the assumption of independence between the datasets used for developing and evaluating our approach.

## Notes

### Competing Interest Statement

The authors have declared no competing interest.

