## Supplemental Information for "Predicting tumour mutational burden from histopathological images using multiscale deep learning"

### Supplementary Methods

#### Multiscale Tiles

Non-overlapping tiles of size 512 × 512 pixels were extracted from each slide at magnifications of 5X, 10X, and 20X. Slides were extracted using the Openslide library[1] (version 3.4.1). The tiles with little tissue present were removed, i.e. tiles with > 50% of their area consisting of background pixels with each RGB value > 220. This process generated approximately one million tiles over the entire dataset.

#### Deep Learning

The neural network loss was weighted for each tile by the reciprocal prevalence of its true class to minimize the impact of class imbalances. Optimization was performed with RMSProp on batches of 22 images, with a learning rate of 1e-2, momentum of 0.95, and epsilon of 1e-10. Training was carried out until convergence, which took between 2 and 15 epochs depending on the model. Our deep learning pipeline was developed using two GeForce GTX 1080 GPUs and the Tensorflow library[2] (version 1.15). Our codebase was written entirely in Python[3] (version 2.7).

#### Random Forest

The random forest classifier had 64 tree estimators and a depth of 4. A number of alternative classifiers in place of the random forest were cross-validated on the validation dataset. This included logistic regression, naive Bayes, k-nearest neighbors, support vector machines, decision trees, latent Dirichlet allocation, quadratic discriminant analysis, and neural networks. Random forest was chosen because it achieved the best receiver operating characteristics on the validation dataset.

**Model Prediction**

It took approximately 40 ms to calculate the probability of each tile and less than 1 min to run our entire pipeline on a single patient using two GeForce GTX 1080 GPUs. It took approximately 2–2.5 min to scan histopathological slides in the same manner as was used for TCGA, using an Aperio scanner. With new ultra-fast digital scanners, it is likely that this step would no longer be a bottleneck.

### Supplementary Figures:

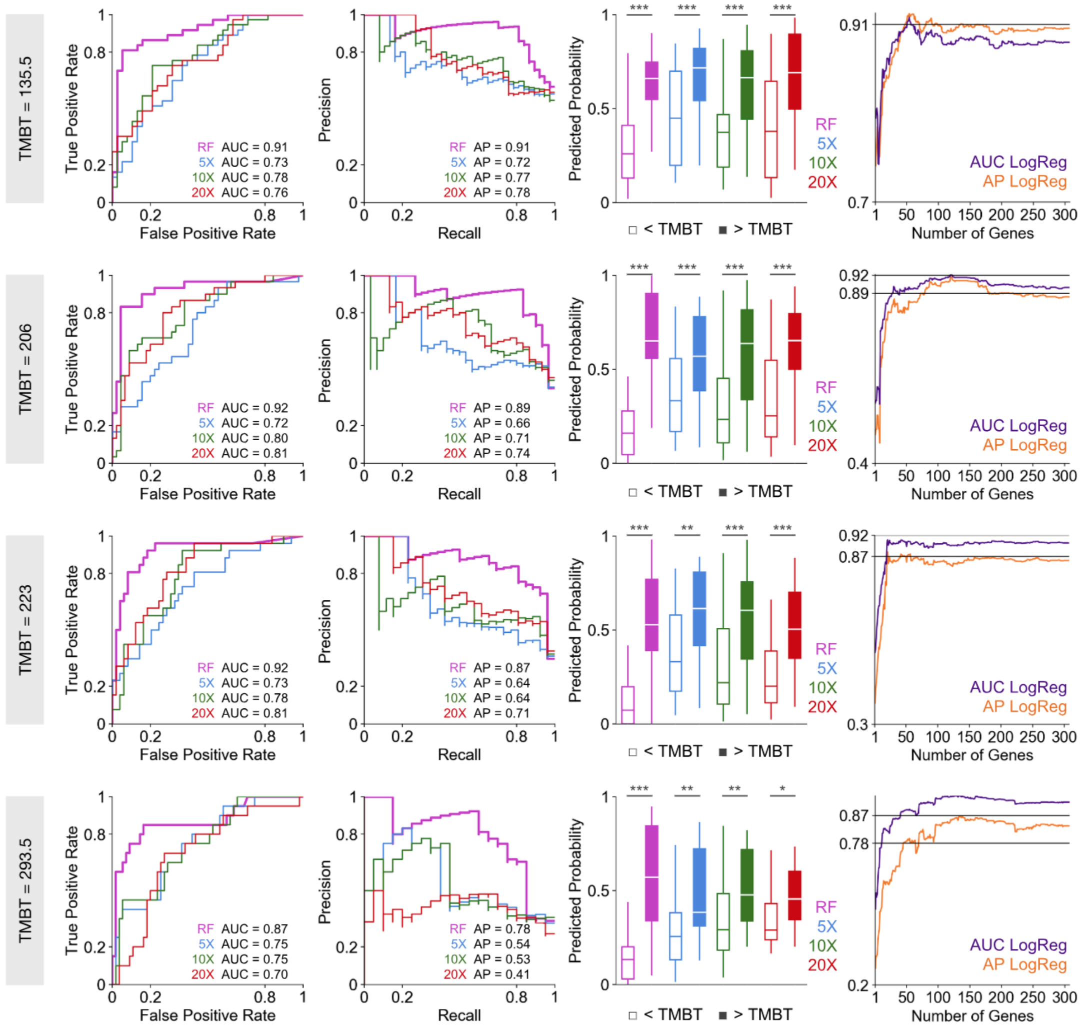


### Supplementary Figure 1.

**Prediction performance at four different TMB thresholds (TMBTs).**

Receiver operator curves (ROC), precision recall curves, and box plots of the probability predictions of Image2TMB and the predictions at each magnification. The area under the ROC curve (AUC) and the area under the precision recall curve (AP) of Image2TMB was compared against a baseline model consisting of a logistic regressor (LogReg) over a selected panel of genes. Confidence intervals are in Supplementary Table 1. All results are reported only for patients in the test dataset.

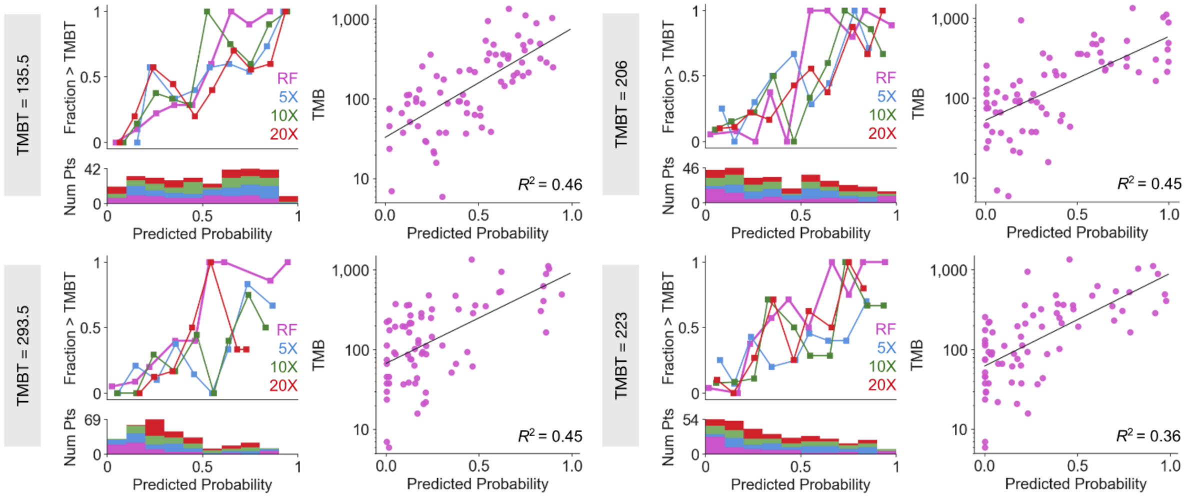


**Supplementary Figure 2.**

**Calibration and regression at the four TMB thresholds (TMBTs).**

Calibration plots for the probability predictions of Image2TMB and the predictions at each magnification. In each scatter plot, each dot corresponds to one patient (Pt), the x-axis is the probability of having high TMB that Image2TMB assigns to the individual and the y-axis is the observed TMB. All results are reported for patients in the test dataset.

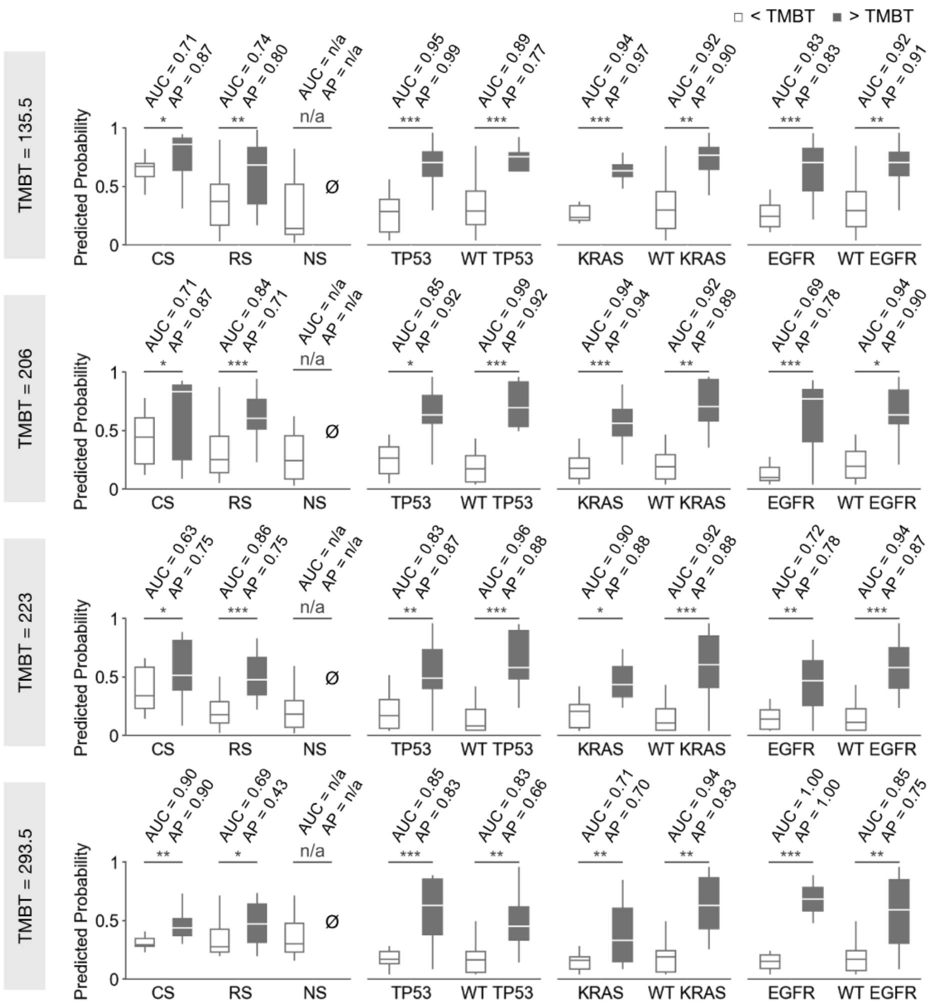


**Supplementary Figure 3.**

**Stratification study of TMB prediction at four TMB thresholds (TMBTs).**

AUC and AP are reported in patient populations stratified by smoking history: current smoker (CS), reformed smoker (RS), nonsmoker (NS); mutated and wild type (WT) TP53; mutated and wild type KRAS; mutated and wild type EGFR. The TMBs of all nonsmokers were below all the TMB thresholds. All results are reported only for patients in the test dataset.

| | RF | | 5X | | 10X | | 20X | |
|---|---|---|---|---|---|---|---|---|
| Threshold | AUC | AP | AUC | AP | AUC | AP | AUC | AP |
| 135.5 | ± 0.037 | ± 0.037 | ± 0.061 | ± 0.062 | ± 0.056 | ± 0.057 | ± 0.058 | ± 0.056 |
| 206 | ± 0.035 | ± 0.041 | ± 0.062 | ± 0.066 | ± 0.054 | ± 0.062 | ± 0.053 | ± 0.060 |
| 223 | ± 0.035 | ± 0.044 | ± 0.061 | ± 0.067 | ± 0.056 | ± 0.067 | ± 0.053 | ± 0.062 |
| 293.5 | ± 0.044 | ± 0.056 | ± 0.059 | ± 0.070 | ± 0.059 | ± 0.070 | ± 0.063 | ± 0.069 |

**Supplementary Table 1.**

**Confidence intervals (95%) of AUC and AP for all thresholds, magnifications, and RF.**

All results are reported only for patients in the test dataset (n=75).

| Threshold | RF | 5X | 10X | 20X |
|---|---|---|---|---|
| 135.5 | 0.62 | 0.53 | 0.56 | 0.55 |
| 206 | 0.62 | 0.53 | 0.58 | 0.56 |
| 223 | 0.64 | 0.54 | 0.57 | 0.55 |
| 293.5 | 0.61 | 0.55 | 0.55 | 0.51 |

**Supplementary Table 2.**

**Matthew’s Correlation Coefficient (MCC) for all thresholds, magnifications, and RF.**

All results are reported only for patients in the test dataset (n=75).
